# Glabridin induces paraptosis-like cell death via ER stress in breast cancer cells

**DOI:** 10.1101/2022.08.02.502466

**Authors:** Xiang Cui, Min Cui

## Abstract

Glabridin, a polyphenolic flavonoid isolated from the root of the *glycyrrhiza glabra*, has been demonstrated to have anti-tumor properties in human malignancies. This study found that glabridin decreased the viability of human breast cancer MDA-MB-231 and MCF7 cells in a dose-dependent manner that was not involved in the caspase-3 cascade. Glabridin promoted the formation of extensive cytoplasmic vacuolation by increasing the expression of endoplasmic reticulum (ER) stress markers BiP, XBP1s, and CHOP. The transmission electron microscopy and fluorescence with the ER chaperon KDEL suggested that the vacuoles were derived from ER. Glabridin-induced vacuolation was blocked when protein synthesis was inhibited by cycloheximide, demonstrating that protein synthesis is crucial for this process. Furthermore, we determined that glabridin causes loss of mitochondrial membrane potential as well as the production of reactive oxygen species, both of which lead to mitochondrial dysfunction. These features are consistent with a kind of programmed cell death described as paraptosis. This work reports for the first time that glabridin could induce paraptosis-like cell death, which may give new therapeutic approaches for apoptosis-resistant breast cancers.

## 1. Introduction

Breast cancer is the most frequently diagnosed cancer and the second-highest cause of cancer-related mortality in women globally [1]. The treatment of breast cancer depends on several factors and can include surgical resection, chemotherapy, radiation, hormonal therapy, or biologic therapy [2]. However, these therapies are unsatisfactory due to metastasis and high recurrence rates [3]. Therefore, it is critical to identify novel therapies that could increase the overall survival of patients with breast cancer.

The majority of the anti-cancer therapies activate apoptosis and related cell death pathways to kill malignant cells [4]. However, cancer cells may evade apoptosis and resist cell death by activating anti-apoptotic and cell survival signals [3, 5]. Hence, the identification of alternative cell death mechanisms may contribute to overcoming apoptosis resistance. Paraptosis is a kind of caspase-independent programmed cell death presenting vacuolation of cytoplasm, swelling of endoplasmic reticulum, and/or mitochondria [6-7]. This form of cell death lacks the morphological characteristics of apoptosis, such as the production of apoptotic bodies, chromatin condensation, and nuclear fragmentation [6-7]. However, the molecular mechanism of paraptosis has not yet been fully understood. Recent studies show that paraptosis is correlated to the activation of mitogen-activated protein kinases (MAPKs) and could be suppressed by cycloheximide, which inhibits translation elongation through binding to the 60S ribosome [8-9]. It is also related to proteostasis disruption and endoplasmic reticulum stress, a perturbation of the ER homeostasis caused by the accumulation of unfolded or misfolded proteins within ER lumen [6]. In addition, disturbances of intracellular calcium homeostasis and the generation of reactive oxygen species (ROS) have been demonstrated to be involved in paraptosis [10].

Multifarious natural and synthetic compounds have been found to induce paraptosis in cancer cells [11-15]. Glabridin (GLA) is a polyphenolic flavonoid originally isolated from the root of *glycyrrhiza glabra* (licorice) [16]. It has shown to be associated with multiple biological properties such as anti-oxidant [17], anti-bacterial [18], anti-inflammatory [19], neuroprotective [20], anti-atherosclerotic [21], and immunomodulatory activities [22]. Previous studies have revealed that glabridin exhibits anti-tumor effects by inhibiting proliferation, migration, and invasion in various human cancers, including breast cancer [23-27]. However, the molecular mechanism behind the anti-tumor effects of glabridin in breast cancer is still unclear.

The purpose of this study is to explore the modalities and mechanism of glabridin induced cell death in human breast cancer cells. We used electron and confocal microscopy, molecular biology, and biochemical approaches to evaluate the morphological and functional responses of breast cancer cells to glabridin treatment.

## 2. Materials and methods

### 2.1. Cell culture and reagents

MDA-MB-231 and MCF7 cells were grown in Dulbecco’s Modified Eagle’s Medium (Gibco, NY, USA) containing 10% fetal calf serum (Biological Industries, Beit HaEmek, Israel). Glabridin (B20474) was purchased from Yuanye Bio-Technology (Shanghai, China). Paclitaxel (P106869), cycloheximide (C112766), and doxorubicin (D107159) were from Aladdin Biochemical Technology (Shanghai, China). GSK2656157 (HY13820) and 4μ8c (HY19707) were from Med-ChemExpress (NJ, USA). MG132 (GC10383) was purchased from Abcam (MA, USA) and Z-VAD-FMK (A1902) was from APExBIO (TX, USA).

### 2.2. Cell viability assay

The cell counting kit-8 (CCK-8; K1018, APExBIO) assay was used to evaluate the cytotoxicity of glabridin. In brief, the cells were plated at 1×10^4^/well in 96-well plates and exposed to 20, 40, 60, 80, and 100μM glabridin for 24 h or 100μM glabridin for 0.5 h, 1 h, 2 h, 6 h, and 24 h. After that, 10 μl CCK-8 reagent was added directly and incubated for another 2 h at 37°C, and then optical density (OD) was measured at 450 nm with a microplate reader (TECAN Infinite M Nano). Data were analyzed using Graph-Pad Prism to derive the EC50.

### 2.3. Flow cytometry analysis

For cell cycle analysis MDA-MB-231 and MCF7 cells (5×10^5^ cells/dish) were seeded in a 60 mm dish and exposed to 100 μM glabridin or 1 μM paclitaxel for 24 h. The cells were then harvested, washed with 1xPBS, and fixed in 70% ethanol at −20°C for 2 h. The fixed cells were centrifuged and washed with 1xPBS two times. One unit of RNase A was added to the cell suspension and incubated for 30 min at 37°C. Subsequently, the cells were resuspended in 300 μl of PI staining buffer and were incubated at 37°C for 10 min before the test. The cells were then analyzed for their DNA content by flow cytometry (Agilent Novocyte Quanteon). For cell death analysis MDA-MB-231 and MCF7 cells were seeded in 60 mm dishes at a density of 5×10 ^5^ cells/well. After treatment with 60 μM glabridin in the absence or presence of Z-VAD-FMK (20 μM), cells were harvested and washed with 1xPBS, and re-suspended in 1xbinding buffer. The cell suspension was incubated with PI staining buffer for 5 min at 37°C in the dark. The cells were then analyzed with flow cytometry.

### 2.4. Colony formation assay

Cells were plated at a density of 5×10^3^ cells per well into 6-well plates and incubated for 24 h at 37°C. After glabridin treatment at indicated concentrations, the cells were allowed to proliferate in a fresh medium. After 10 days, colonies were fixed with 4% paraformaldehyde for 10 min and stained with 0.1% crystal violet (Solarbio, Beijing, China) for 30 min for colony visualization [13].

### 2.5. Transfection and confocal microscopy

Cells were seeded at a concentration of 1×10^5^ cells per well in a 24-well plate and were transfected with KDEL-GFP or COX8-RFP plasmid using ScreenFect A (WAKO, Japan) according to the manufacturer’s protocol. After 24 h, cells were treated with 100 μM glabridin for another 24 h, and fluorescence and differential interference contrast (DIC) images of cells were visualized under a confocal microscope (Nikon AX).

### 2.6. Transmission electron microscopy

Cells were treated with glabridin, and initially fixed with 2.5% glutaraldehyde for 1 h. Then the cells were postfixed in 1% osmium tetroxide for 1 h at 4°C. After dehydration with graded ethanol series, the cells were embedded in EMBED 812, polymerized, and observed under the electron microscope (HT7800, HITACHI).

### 2.7. RT-PCR

Total RNA was extracted from cells using TRNzol reagent (TIANGEN, China) according to the manufacturer’s protocol. First-strand cDNA was synthesized in a 20 μl of reaction volume using a random primer (TAKARA, Japan) and 1 μl reverse transcription enzyme M-MLV-RT (Promega, USA). PCR was performed using each specific primer set in a total volume of 20 μl containing 10 pmol of each primer, 4 μmol dNTPs (Solarbio, China), 1 unit of Taq DNA polymerase (TIANGEN, China), and 1 × PCR buffer. Primer sequences are summarized in Supplementary Table S1. PCR cycle parameters were 15 s at 94°C, 15 s at 55°C, and 20 s at 72°C for 16 - 25 cycles (BiP, 22; CHOP, 25; XBP1, 25; 28S, 16) followed by 72°C for 5 min. Aliquots (5 μl) of each reaction mixture were electrophoresed on 4.8% polyacrylamide gels.

### 2.8. Western blotting

For western blot analyses, proteins were extracted from cells using cell extraction buffer containing 20 mM Tris-HCl (pH 7.5), 500 mM NaCl, 10 mM MgCl_2_, 2 mM EDTA, 10% glycerol, 1% Triton X-100, 2.5 mM β-glycerophosphate, 1 mM NaF, 1 mM DTT and 1 mM complete protease inhibitors (APExBIO) at 4 °C. After centrifugation, soluble protein in the lysates was quantified. Samples were separated by 8–15% sodium dodecyl sulfate-polyacrylamide gels and transferred to a PVDF membrane (IPVH00010 Millipore Immobilon). After transfer, the membrane was blocked with 5% BSA, washed with TBST, and incubated with primary antibodies overnight at 4 °C. Then, the membrane was washed with TBST, incubated with a secondary antibody for 1 h at room temperature, washed, and detected with a gel imager (Bio-Rad, Hercules, CA, USA). Anti-GAPDH (T0004, 1:3000) and anti-cleaved caspase-3 (AF7022, 1:500) were from Affinity Biosciences (Ohio, USA). Anti-cleaved PARP (T40050s, 1:1000) was from Abmart (Shanghai, China) and Anti-ubiquitin (H1021, 1:1000) was from Santa Cruz Biotechnology (California, USA).

### 2.9. Mitochondrial membrane potential assay

Mitochondria membrane potential is measured by JC-1 (Beyotime Biotechnology, China) staining. Following treatment with glabridin or CCCP (Beyotime Biotechnology, China), the cells were incubated with (10 μg/ml) JC-1 for 30 min at 37°C. Cells were then visualized by a fluorescence microscope. For quantitative analysis, the cells were treated with glabridin and subsequently incubated with JC-1 as described above and were washed with cold staining buffer. Then, the cells were analyzed by a fluorescence microplate reader [INFINITE M NANO, TECAN], and the mitochondrial membrane potential was indicated by the ratio of green/red fluorescence intensity.

### 2.10. Reactive oxygen species assay

Cells were grown on a coverslip in 24-well plates for 24 h and then treated with 100 μM glabridin or 100 ug/ml Rosup for 6 h. The cells were then treated with 10 μM DCFH-DA (Biosharp, China) for 30 min before being imaged using an inverted microscope (Invitrogen EVOS M5000).

### 2.11. Statistical analyses

Data are indicated as mean ± s.d. from at least three independent experiments for each experimental condition. Student’s t-test was performed to calculate P values. P < 0.05 was defined as significant.

## 3. Results

### 3.1. Glabridin treatment inhibits cell viability in breast cancer cells

To elucidate the anti-cancer effect of glabridin, we first investigated its cytotoxicity in various breast cancer cells. Glabridin significantly reduced the viability of breast cancer cells at concentrations ranging from 10 μM to 100 μM (Figure S1), which is comparable to the dosages at which glabridin inhibits other cancer cells [24, 26]. To assess the most effective concentrations of glabridin on breast cancer cells, we treated MDA-MB-231 and MCF7 cells for 24 h with 20 μM, 40 μM, 60 μM, 80 μM, and 100 μM glabridin. As shown in Figure 1A, glabridin treatment significantly reduced the viability of MDA-MB-231 and MCF7 cells in a dose-dependent manner. The half-maximal effective concentration (EC50) values of glabridin were 42.74 μM for MDA-MB-231 cells and 49.5 μM for MCF-7 cells, respectively. Moreover, at higher glabridin concentrations (100 μM), cell viability was drastically decreased to a level equivalent to that of the 24 h treatment group in the early stages of exposure (0.5 h) (Figure 1B). Flow cytometry was then used to study the effects of glabridin on cell cycle distribution. Glabridin treatment caused cell cycle arrest in both MDA-MB-231 and MCF7 cells at the G0/G1 and S phases (Figure 1C). This is in contrast to the G2/M cycle arrest caused by paclitaxel, a major breast cancer chemotherapy drug known to trigger apoptosis [28] (Figure 1C). To further estimate the long-term cytotoxic effect of glabridin, we evaluated colony formation capacity in breast cancer cells. Following 10 days of glabridin treatment, the colony-forming ability of the cells drastically reduced when the concentration of glabridin was above 40 μM (Figure 1D).

**Figure 1.**
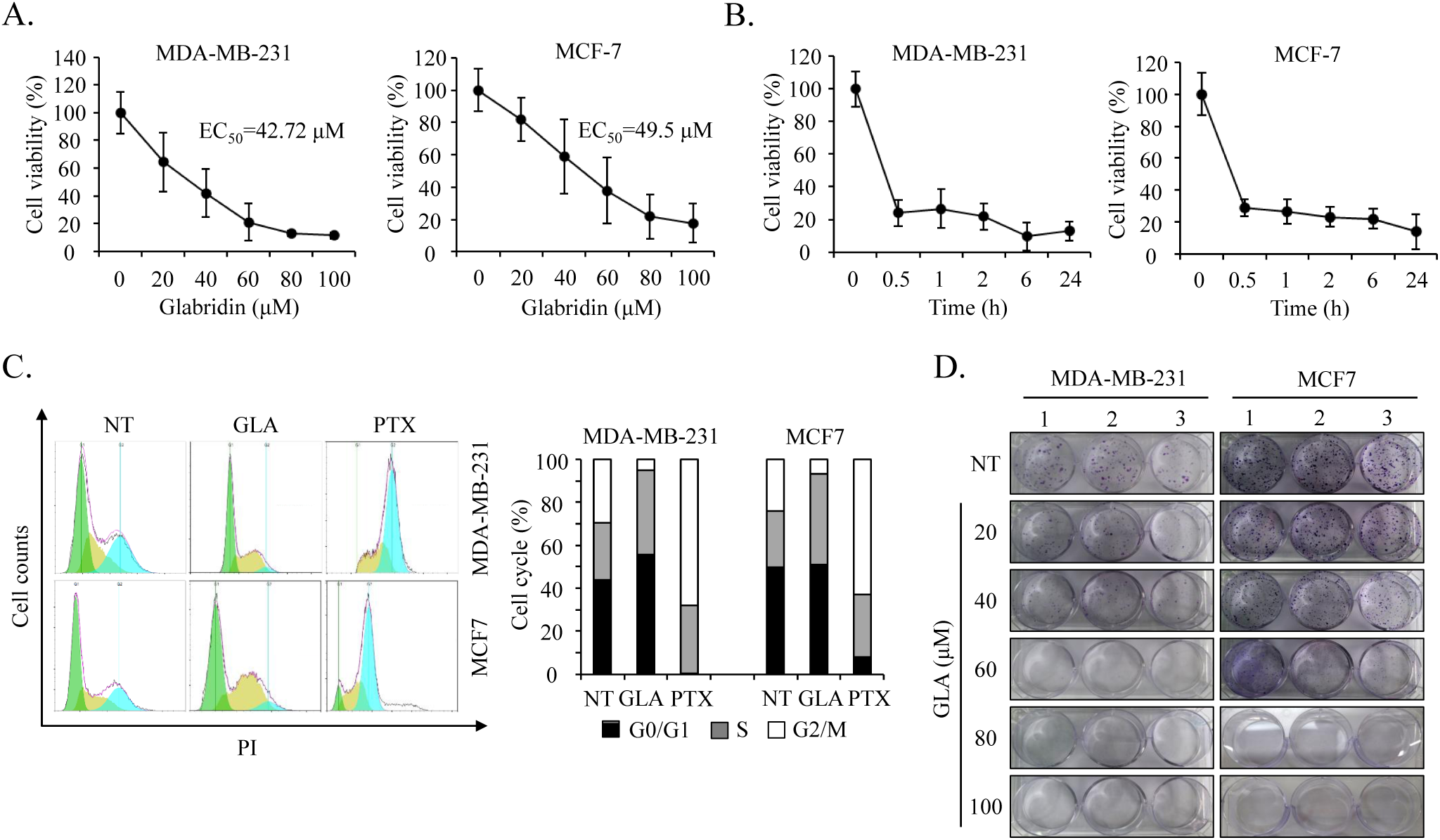
Glabridin inhibits cell viability, clonogenic capacity, and cell cycle progression of breast cancer cells. **(A) and (B)** The cytotoxicity of glabridin against MDA-MB-231 and MCF7 cells was assessed by CCK-8 assays. The cells were treated with (A) indicated doses of glabridin for 24 h or (B) 100 μM glabridin for indicated time periods. The OD values were measured in a microplate reader and the EC50 values of glabridin were calculated. The results are presented as the means ± SD of four independent experiments. **(C)** Flow cytometry histograms (left panel) and quantification (right panel) of PI-labeled cells in each phase of the cell cycle. Cells were treated with 100 μM glabridin or 1 μM paclitaxel for 24 h and were then stained with PI for flow cytometric analysis. **(D)** Colony formation abilities of MDA-MB-231 and MCF7 cells treated with increasing concentrations of glabridin for 10 days. GLA, glabridin; NT, no treatment; PTX, paclitaxel; PI, propidium iodide

### 3.2. Glabridin induces cytoplasmic vacuolation in a concentration-dependent manner

We next evaluated the morphological changes in MDA-MB-231, MCF7, MDA-MB-435, and MDA-MB-468 cells after treatment with glabridin. Interestingly, extensive cytoplasmic vacuolation was observed in all four breast cancer cell lines (Figure 2A and Figure S2). These vacuoles resemble those produced by honokiol [11] or curcumin [12], both of which have been shown to induce paraptosis (Figures S3 and S4). Moreover, the number of vacuoles grows and fuses together as the concentration of Glabridin increases (Figure 2A). With high glabridin concentrations (100 μM), small vacuoles developed within 0.5 h of stimulation and grew in number and size as stimulation time increased until they coalesced (Figure 2B).

**Figure 2.**
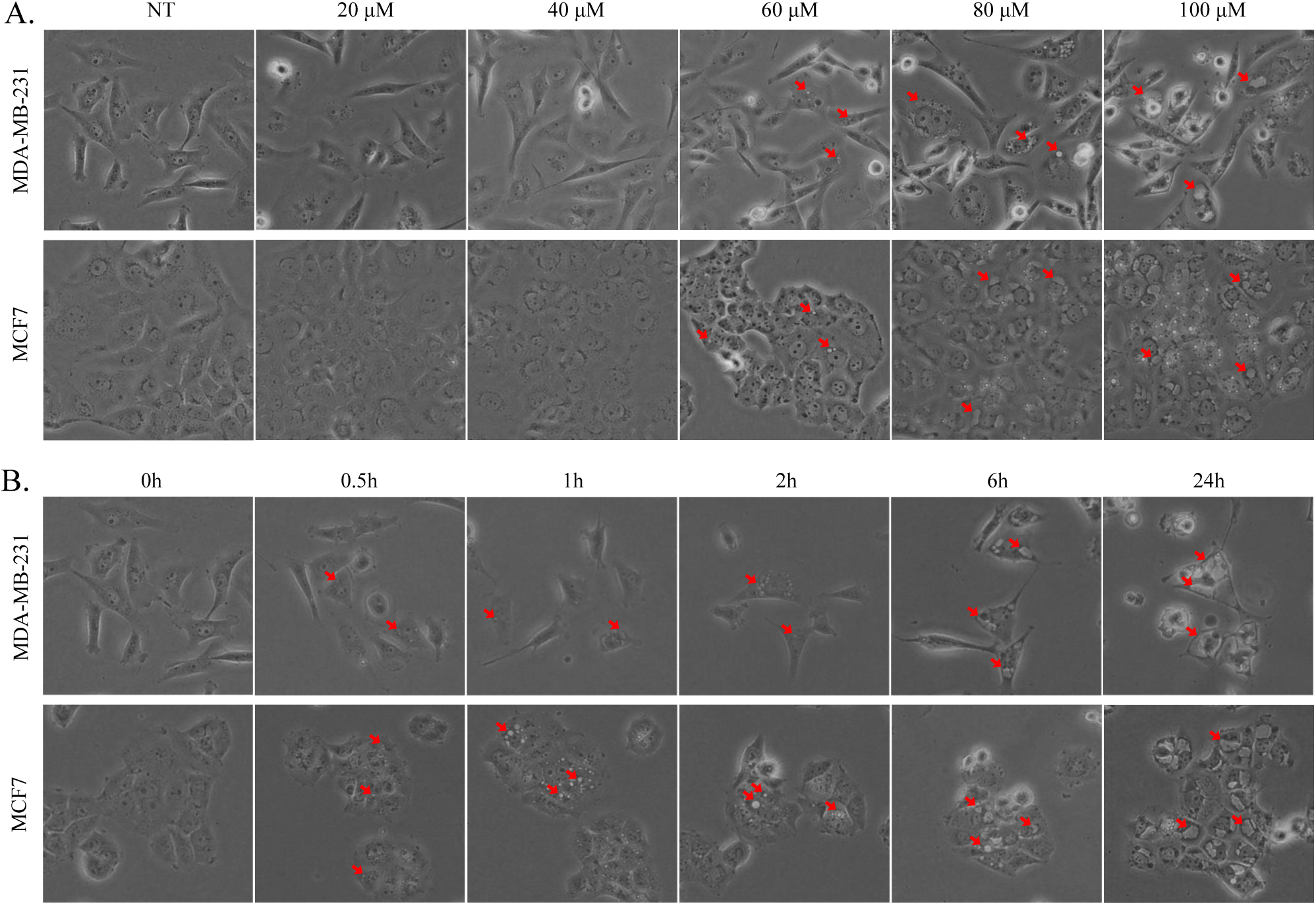
Glabridin induces cytoplasmic vacuolation in breast cancer cells. **(A) and (B)** Morphological changes of MDA-MB-231 and MCF7 cells treated with glabridin observed under an inverted light microscope. The cells were treated with (A) increasing concentrations of glabridin for 24 h or (B) 100 μM glabridin for indicated time periods. Cytoplasmic vacuoles were indicated by the red arrows.

### 3.3. Glabridin induces ER-derived vacuolation due to ER stress and proteasome inhibition

To further evaluate the origin of cytoplasmic vacuoles in glabridin-treated cells, we expressed endoplasmic reticulum (ER) marker KDEL and mitochondrial inner membrane marker COX8 in glabridin-treated MDA-MB-231 and MCF7 cells. As expected, ER in control cells exhibited a reticular pattern, whereas mitochondria in control cells showed elongated and filamentous morphologies (Figure 3A). In contrast, cells treated with glabridin distributed dilated ER and mitochondria (Figure 3A, Video S1, and S2). Moreover, the vacuolar membrane in the glabridin treated cells appeared positive for ER marker suggesting that ER is one potential source of membranes for vacuoles. In addition, our time-lapse imaging revealed that glabridin first caused mitochondrial swelling, and then fused into oval or spherical mitochondria (Video S1). Electron microscopy further revealed that large empty vacuoles surrounded the intact nuclei and the ER and mitochondria exhibited swelling in the cells treated with glabridin (Figure 3B).

**Figure 3.**
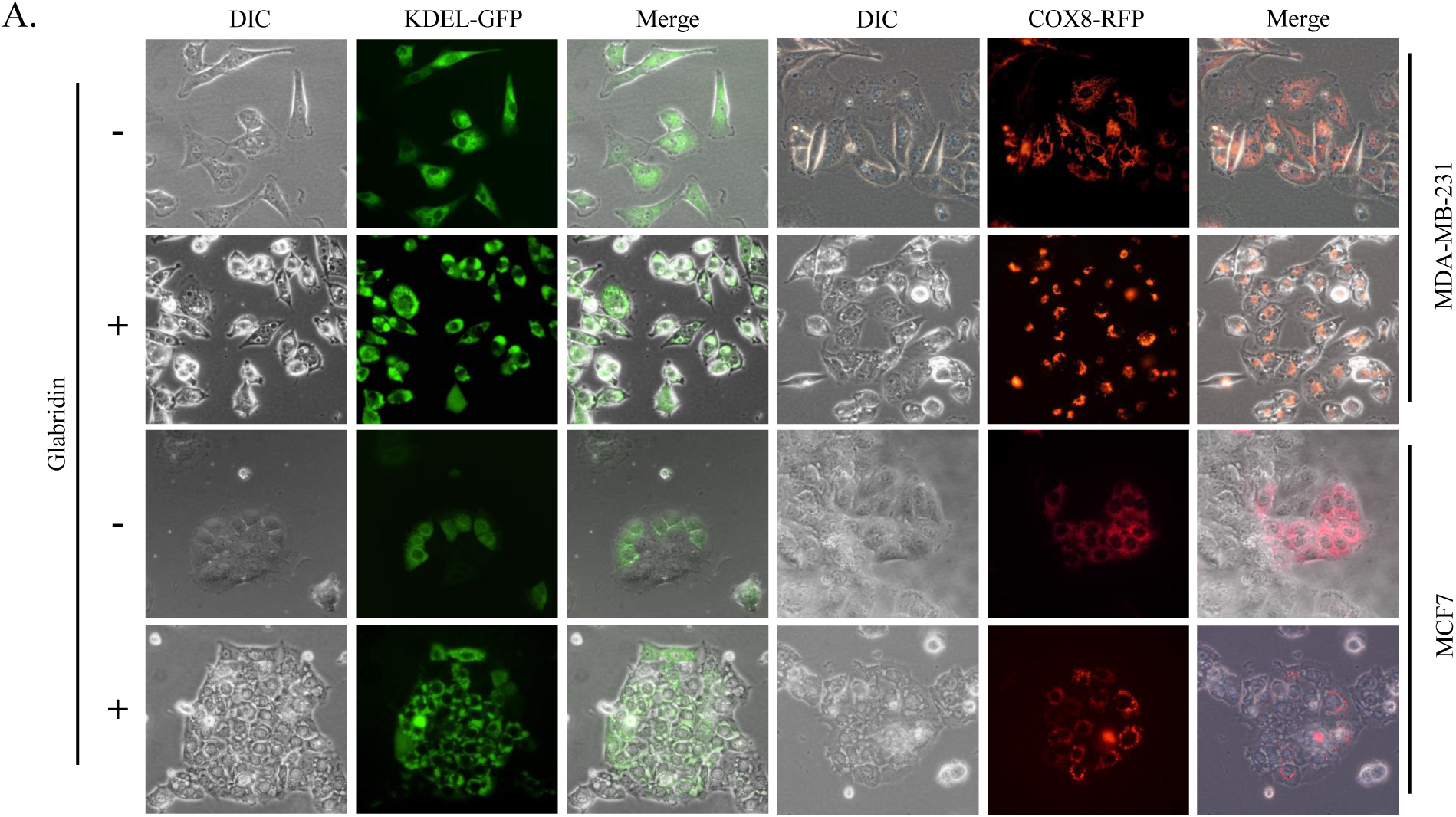
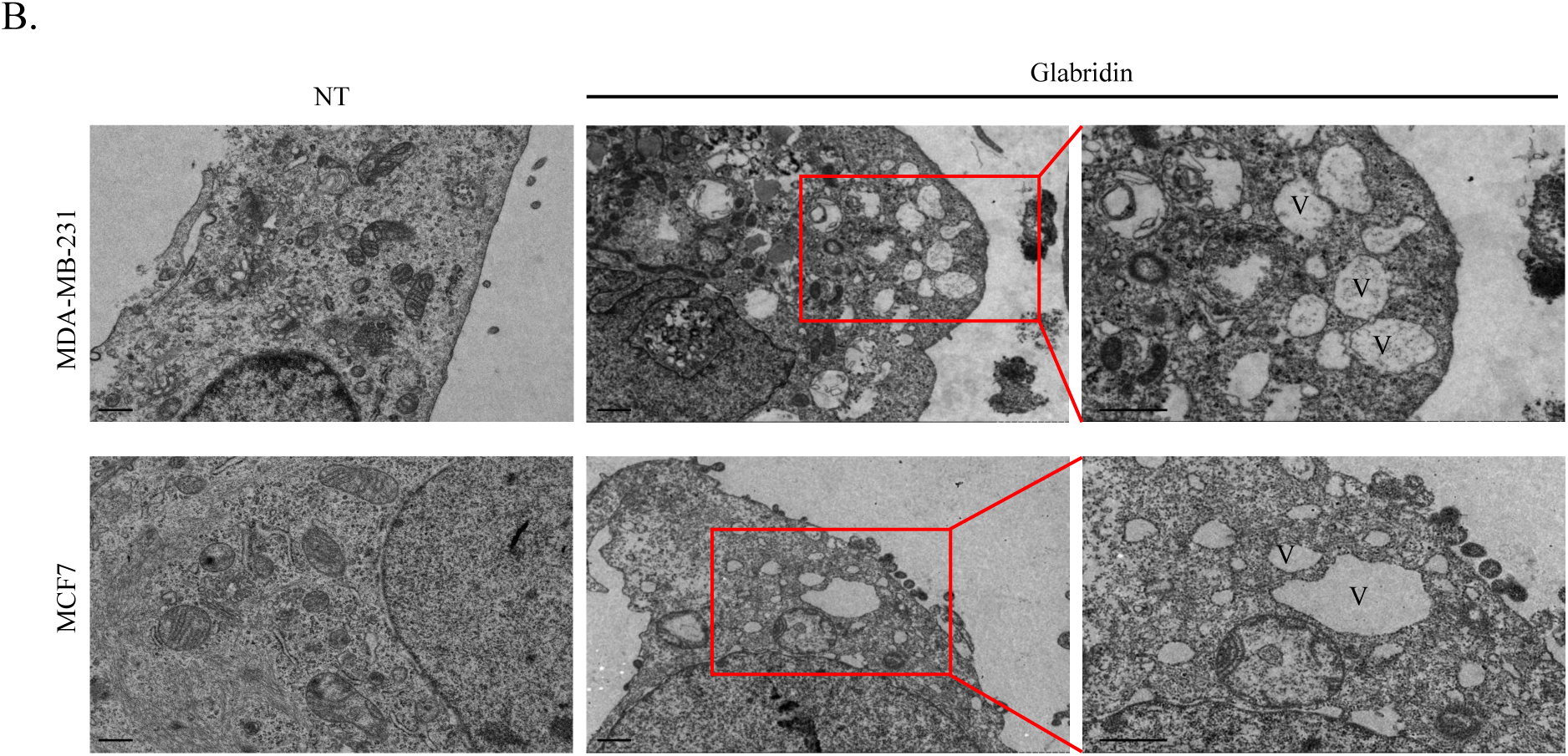
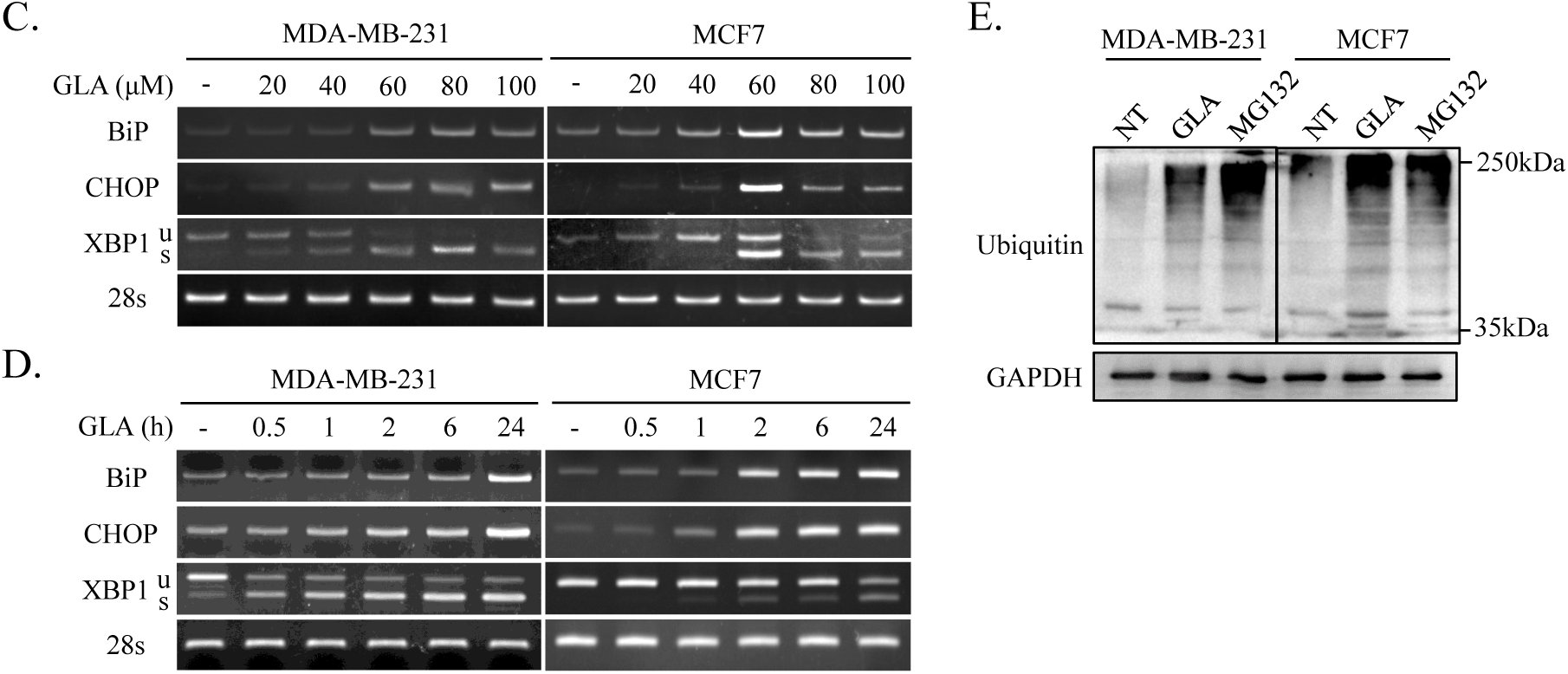
Glabridin induces ER and mitochondrial dilation, ER stress, and proteasome inhibition. **(A)** Representative fluorescence images of MDA-MB-231 and MCF7 cells after glabridin treatment. Cells were transiently transfected with the KDEL-GFP or COX8-RFP plasmids 24 h before treatment with glabridin (100 μM). After 24 h, the cells were observed under fluorescence microscopy. **(B)** Transmission electron micrograph images of cells treated with glabridin (100 μM) for 6 h. The inset shows a part of a high magnification image. Scale bars, 1 μm; V, vacuole; NT, no treatment. **(C) and (D)** The mRNA expression of ER stress markers in cells treated with glabridin. The cells were treated with (C) indicated concentrations of glabridin for 24 h or (D) 100 μM glabridin for indicated time periods. 28s was used as a loading control in RT-PCR. **(E)** Cells treated with 100 μM glabridin or 100nM MG132 for 24 h were subjected to western blot analysis with anti-ubiquitin or anti-GAPDH as indicated. The results are presented as the means ± SD of three independent experiments (vs. control: *P<0.05). GLA, glabridin; XBP1u, unspliced XBP1; XBP1s, spliced XBP1; NT, no treatment

It was reported that proteasomal dysfunction and ER stress regulate paraptotic cell death in cancer cells [6]. Next, we examined whether glabridin causes ER stress. RT-PCR analysis revealed that glabridin treatment increased the mRNA expression levels of glucose-regulated protein (BiP/GRP78), spliced X-box binding protein 1 (XBP1s), and C/EBP homologous protein (CHOP) in a dose-and time-dependent manner (Figure 3C and 3D). To establish whether the ubiquitination occurs in the glabridin-treated cells, we examined the expression levels of ubiquitinated protein by western blot analysis. As shown in figure 3E, glabiridin treatment notably upregulated the expression of polyubiquitinated proteins and the ubiquitination level was on par with that of cells treated with MG132, a well-known proteasomal inhibitor. Taken together, these results suggest that glabridin induces ER-derived vacuolation due to ER stress and proteasome inhibition.

### 3.4. Glabridin induces caspase-independent cell death in breast cancer cells

We next treated MDA-MB-231 and MCF7 cells for 24 h with 100 μM glabridin to get insight into the cell death process. Western blotting revealed that the apoptotic markers cleaved PARP and caspase-3 were marginally increased in glabridin-treated MCF7 cells but not in MDA-MB-231 cells (Figure 4A). DAPI staining was further used to examine the effect of glabridin on nuclear changes. Glabridin-treated cells displayed greater fluorescence intensity than control cells, as seen in Figure 4B, due to chromatin condensation. Consistently, the DNA fragmentation assay also showed slight fragmentation of the DNA when the cells were exposed to glabridin (Figure 4C). To investigate the role of caspases in glabridin-induced cytotoxicity, MDA-MB-231 and MCF7 cells were treated with Z-VAD-FMK (a pan-caspase inhibitor), and the fraction of live cells and PI-positive cells was determined by flow cytometry (Figure 4D). The proportion of live cells in the glabridin treatment group (83.34% and 50.62%) was substantially lower than in the control (94.33% and 87.18%) groups. Pretreatment with Z-VAD-FMK, on the other hand, did not protect cells against glabridin-induced cell death. Meanwhile, the CCK-8 assay revealed that Z-VAD-FMK did not reverse the inhibitory effect of glabridin on cell survival (Figure 4E). These findings support the idea that the cell death caused by glabridin in breast cancer cells is triggered by a mechanism independent of caspase family protease activation.

**Figure 4.**
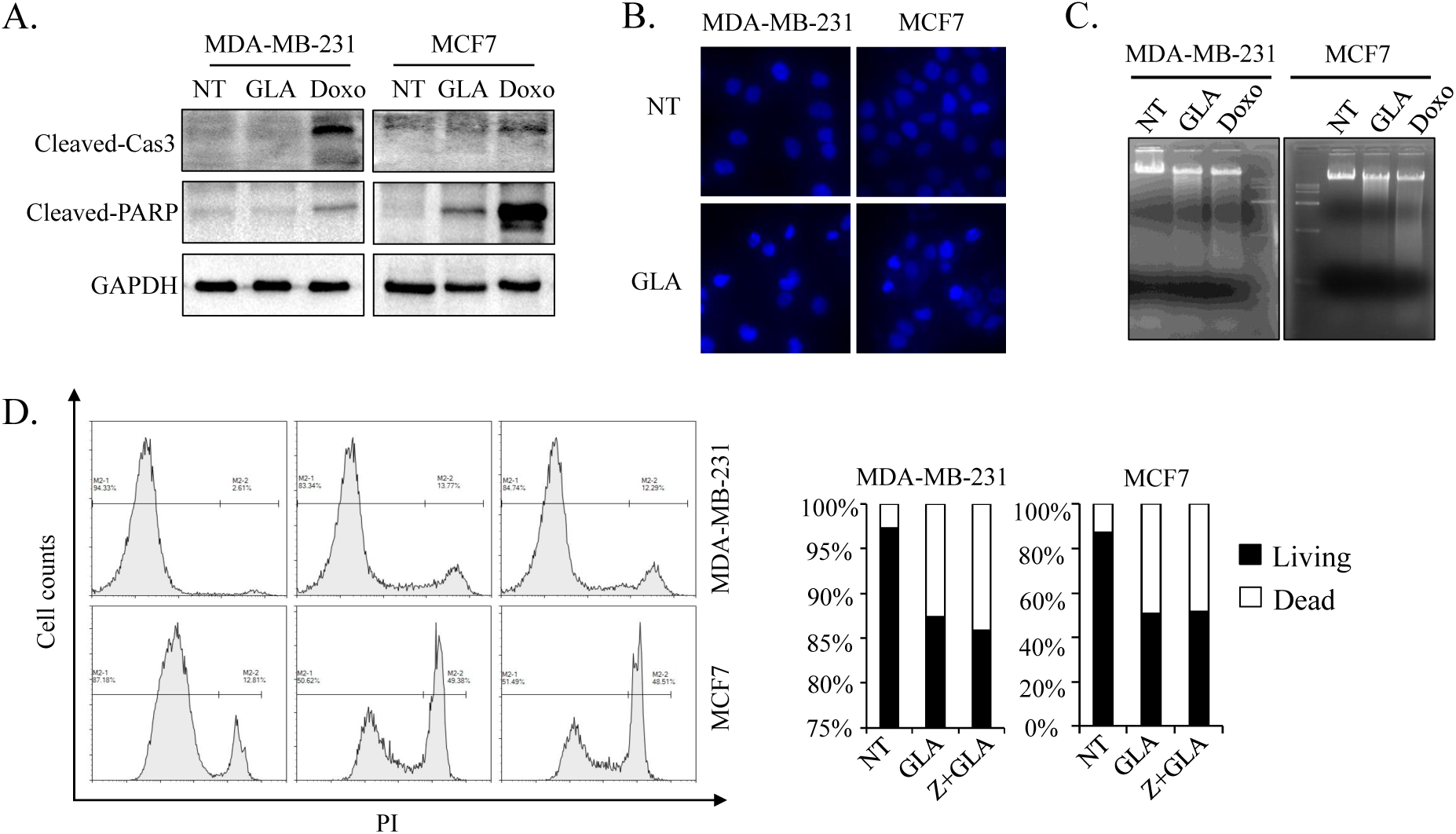
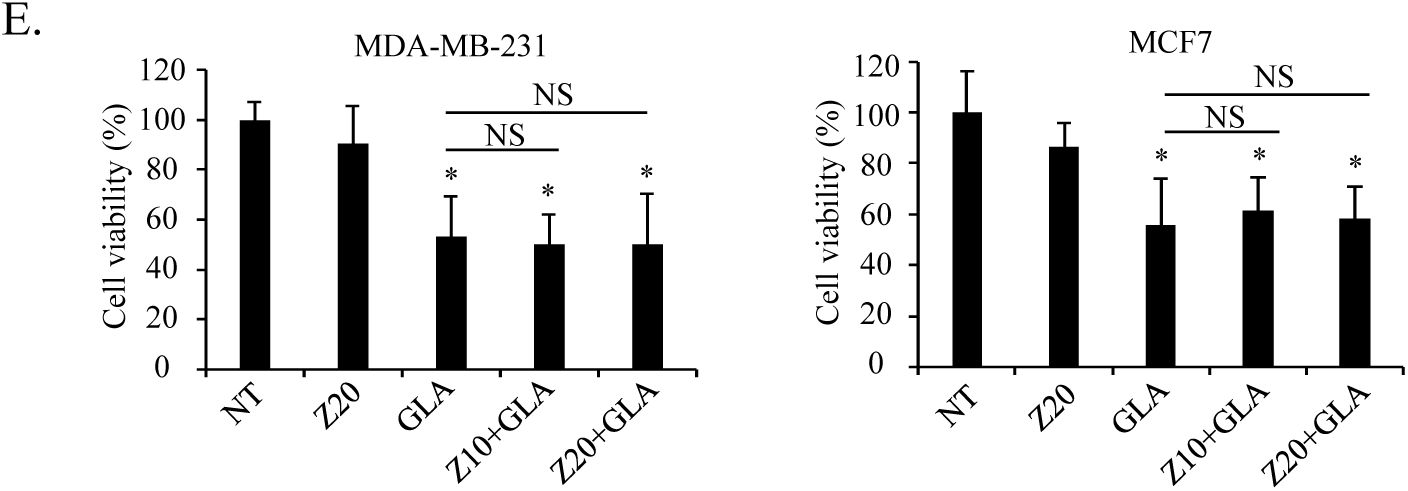
Glabridin induces caspase-independent cell death in breast cancer cells. **(A)** Western blot analysis of caspase-3 and PARP expression in 100 μM glabridin-or 1 μg/ml doxorubicin-treated cells. **(B)** Representative images of nuclear staining. MDA-MB-231 and MCF7 cells were treated with 100 μM glabridin for 24 h, and the nuclei were stained with DAPI. **(C)** DNA fragmentation assay. Genomic DNA isolated from glabridin-or doxorubicin-treated cells subjected to agarose gel electrophoresis. **(D)** Cells were treated with 60 μM glabridin in the absence or presence of Z -VAD-FMK (20 μM), and the percentage of cell death was determined using PI staining. **(E)** Cells were treated with 60 μM glabridin in the absence or presence of Z -VAD-FMK (10 μM or 20 μM), and the cell viability was determined using CCK-8 assay. The results are presented as the means ± SD of three independent experiments (vs. control: *P<0.05; NS, not significant). NT, no treatment; GLA, glabridin; Doxo, doxorubicin; Z, Z-VAD-FMK; Z10, 10μM Z-VAD-FMK; Z20, 20μM Z-VAD-FMK

### 3.5. Glabridin promotes ROS generation and mitochondrial membrane potential loss in breast cancer cells

Mitochondria play an important role in cell death, and mitochondrial membrane potential is a key component that influences mitochondrial activity [29]. To assess changes in mitochondrial membrane potential following glabridin treatment, we employed fluorescent JC-1 labeling. In healthy cells with normal MMP, JC-1 forms red fluorescent aggregates [30]. In contrast, in sick cells with low MMP, JC-1 stays monomeric, displaying only green fluorescence [30]. Following glabridin treatment, the intensity of green fluorescence increased while the intensity of red fluorescence dropped (Figure 5A). Correspondingly, quantitative analysis revealed that glabridin-treated cells had a higher green/red fluorescence intensity ratio when compared to controls, showing that glabridin promotes MMP loss in breast cancer cells (Figure 5B). Moreover, we used 2’,7’–dichlorofluorescein diacetate (DCFH-DA) staining to assess ROS production in glabridin-treated cells. DCFH-DA is hydrolysis and subsequently oxidized to a fluorescent state by ROS, giving a measurement of ROS levels in cells. Microscopic analysis with DCFH-DA revealed that glabridin increases ROS levels in MDA-MB-231 and MCF7 cells (Figure 5C). These data imply that glabridin can promote mitochondrial dysfunction in breast cancer cells.

**Figure 5.**
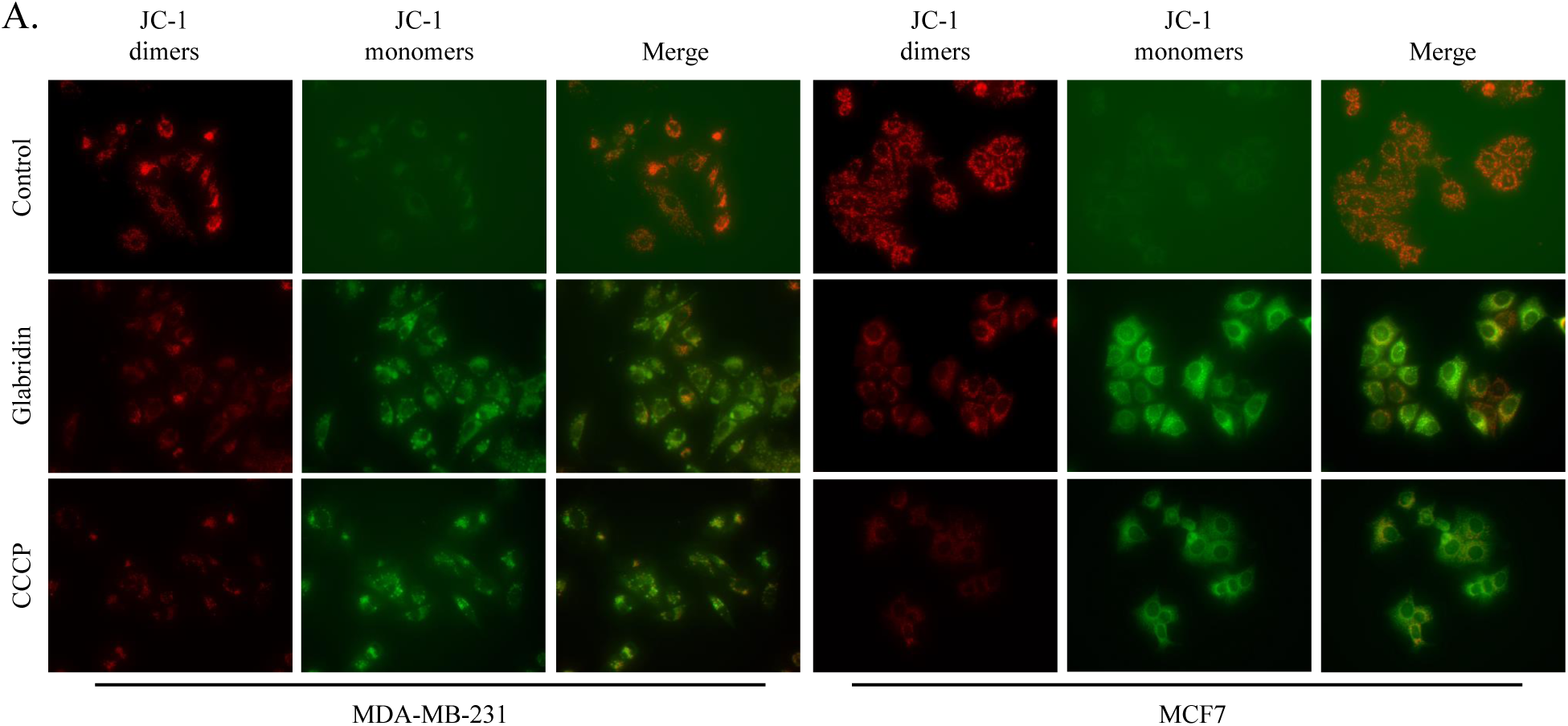
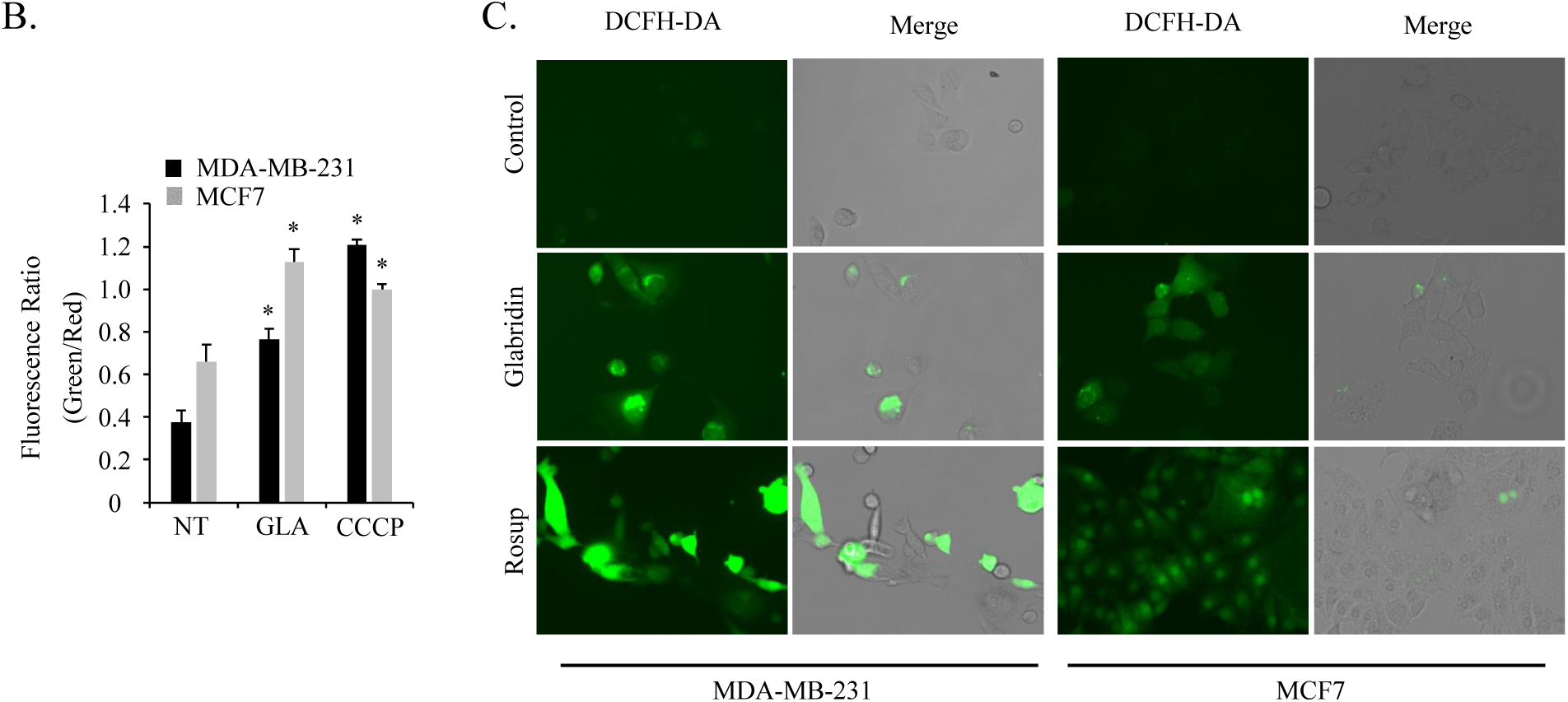
Glabridin induces caspase-independent cell death in breast cancer cells. **(A) and (B)** Mitochondrial membrane potential (MMP) of the glabridin-treated cells was measured by (A) JC-1 fluorescence staining and (B) a fluorescence microplate reader. The graph represents the ratio of green (monomers) and red (dimers) fluorescence. CCCP treatment was used as a positive control. **(C)** Fluorescence images of cellular ROS production in untreated or glabridin or Rosup treated cells. The results are presented as the means ± SD of three independent experiments (vs. control: *P < 0.05). GLA, glabridin; NT, no treatment

### 3.6. Inhibition of PERK synergistically enhances glabridin-induced cytoplasmic vacuolation

ER stress triggers an adaptive signaling network known as the unfolded protein response (UPR), which improves cell survival by lowering unfolded/misfolded protein levels [31]. To investigate the relationship between glabridin-induced paraptosis and the UPR signaling pathway, we pretreated the cells with either the PERK inhibitor GSK2656157 (GSK) or the IRE1 inhibitor 4μ8C before glabridin treatment. As expected, treatment with GSK and 4μ8C inhibited glabridin-induced upregulation of CHOP and spliced XBP-1 (XBP1s) (Figure 6A). On the contrary, proteasome inhibitor MG132 increased mRNA levels of BiP and XBP1s (Figure 6A). Moreover, the addition of cycloheximide (CHX), a protein synthesis inhibitor that has been reported to suppress paraptosis [8], reduced the expression levels of the ER stress markers and cytoplasmic vacuolation in glabridin-treated cells (Figure 6A and Figure S5). Phase-contrast microscopy showed that pre-treatment with MG132 enhanced the glabridin-induced cytoplasmic vacuolation (Figure 6B). Interestingly, despite the fact that GSK alone did not form vacuoles, co-treatment with GSK and glabridin promoted cytoplasmic vacuolation (Figure 6B and Figure S6). To further analyze the effect of GSK on glabridin-mediated cytotoxicity and mitochondrial dysfunction, we performed CCK-8 and mitochondrial membrane potential assay in cells co-treated with glabridin and other drugs as described above. The result revealed that glabridin treatment led to a considerable decrease in cell activity (Figure 6C). However, we did not observe any dramatic changes when the cells co-treated glabridin with GSK or other drugs (Figure 6C and 6D). These results suggest that inhibition of PERK synergistically enhances glabridin-induced cytoplasmic vacuolation, and that the decrease in cell activity caused by glabridin is not positively associated with the formation of vacuoles.

**Figure 6.**
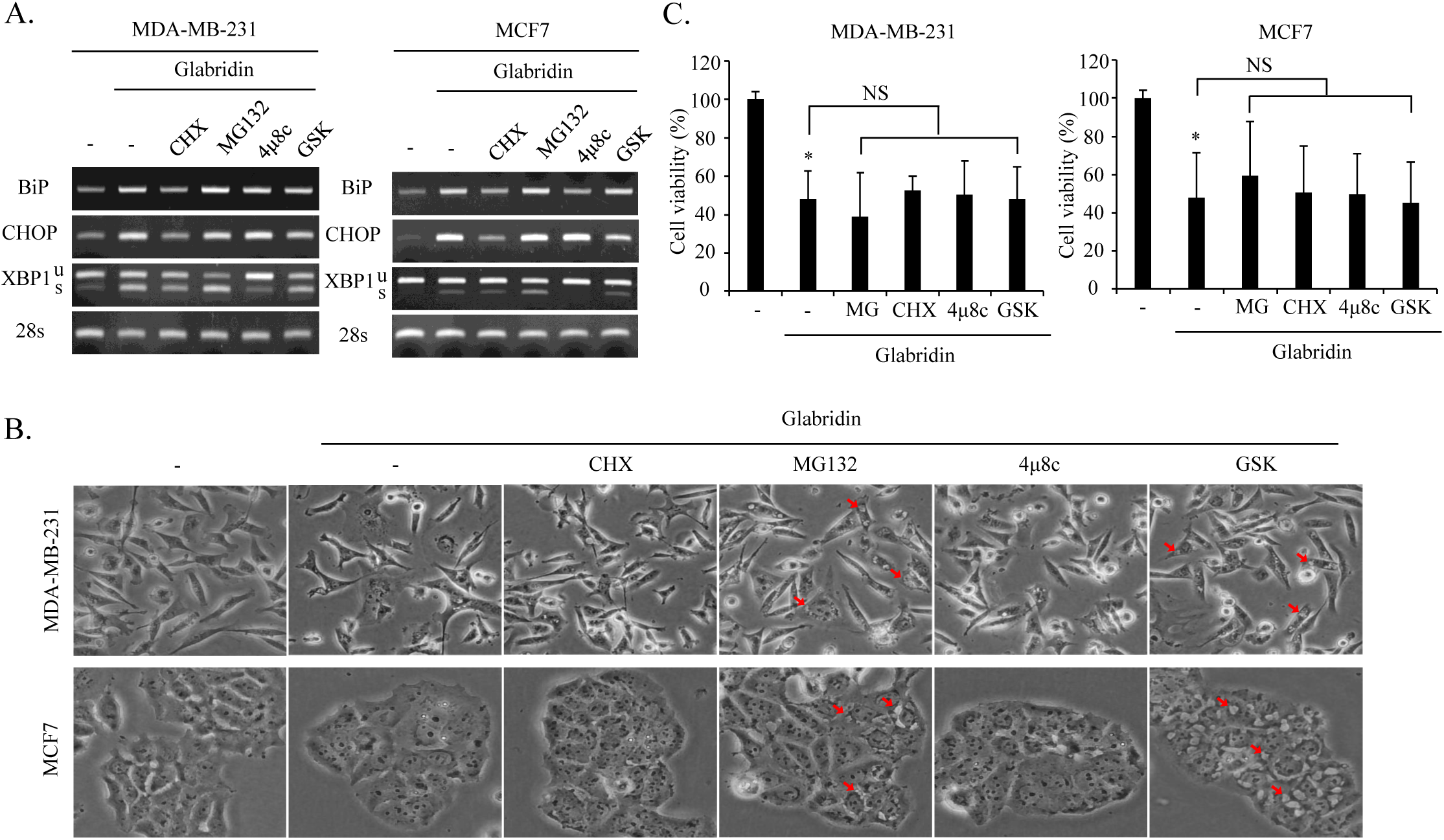
Inhibition of PERK enhances glabridin-induced cytoplasmic vacuolation. **(A)** Cells were pretreated with the indicated stimuli (500 nM CHX; 100 nM MG132; 10 μM 4μ8C; 1 μM GSK) for 20 min prior to the treatment with 60 μM glabridin for 24 h, and then the total RNA was extracted and subjected to RT-PCR. **(B)** Cells were treated as in (A), and the cell viability was determined using CCK-8 assay. The results are presented as the means ± SD of three independent experiments (vs. control: *P < 0.05; NS, not significant). **(C)** Cells were treated as in (A), and observed under the phase-contrast microscope. Cytoplasmic vacuoles were indicated by the red arrow. CHX, cycloheximide; MG, MG132; GSK, GSK2656157

## 4. Discussion

Glabridin is a skin whitening and antioxidant ingredient that is commonly used in cosmetics [32]. Recent studies have shown that glabridin also has growth inhibitory capabilities against a variety of human cancers, including breast cancer [23-27]. Here we show that glabridin treatment induces cytoplasmic vacuolation and caspase-independent paraptosis-like cell death in breast cancer cells.

Glabridin exhibited suppressive effects on cell survival, clonogenic capacity, and cell cycle progression, which was consistent with prior findings [24,27]. Furthermore, we detected extensive cytoplasmic vacuoles in glabridin-treated MDA-MB-231 and MCF7 cells due to ER and mitochondrial dilatation, which is one of the key features of paraptosis. We observed that the vacuoles induced by glabridin grow progressively in size with increasing concentration and duration, eventually fusing into giant vacuoles. Moreover, in line with previous studies [11-12], vacuole formation could be effectively prevented by pretreatment with the protein synthesis inhibitor cycloheximide. Simultaneously, we found that glabridin treatment elevated the expression levels of ER stress markers such as BiP, XBP1s, and CHOP in a time-and concentration-dependent manner, and that this increase happened at the same time as the appearance of cytoplasmic vacuolation. Furthermore, we found that glabridin treatment induced the accumulations of ubiquitinated proteins, indicating that glabridin may prevent protein degradation. While exploring the relationship between UPR signaling and the paraptosis-like cell death induced by glabridin, we noticed that the proteasome inhibitor MG132 and the PERK inhibitor GSK enhanced glabridin-induced cytoplasmic vacuolation. PERK is an ER-resident kinase that phosphorylates the eIF2α under ER stress, which inhibits protein synthesis and restricts the continued influx of ER client proteins [31]. Inhibition of proteasome or PERK leads to an accumulation of unfolded proteins in the ER. However, GSK treatment alone did not trigger vacuole formation, indicating that proteasome dysfunction is required for vacuole formation.

Previous studies have shown that the caspase-dependent apoptotic cascade is crucial for glabridin-induced cell death [24, 26]. Despite the fact that glabridin causes modest caspase-3 and PARP activation in MCF7 cells, the caspase inhibitor Z-VAD-FMK was unable to restore the cytotoxicity caused by glabridin in both MCF7 and MDA-MB-231 cells, demonstrating that apoptosis is not the major cause of glabridin-induced cell death. These differences may be due to the use of different cell types or glabridin concentrations.

Our results reveal that 20 μM glabridin significantly suppressed cell activity, while the vacuoles started to appear when the concentration of glabridin was up to 60 μM. Additionally, we noticed that applying MG132 or GSK, both of which can increase glabridin-induced vacuolation, did not enhance the glabridin-induced reduction in cell activity. Moreover, when we treated cells with a higher concentration of glabridin (100 μM), we saw a significant decrease in cell activity in the early stages (0.5 h) of treatment. Simultaneously, vacuoles began to appear and subsequently grew progressively as the glabridin treatment duration increased. In contrast, the cell activity did not further decrease with the increase in treatment time. Furthermore, our time-lapse images revealed that the morphological alterations in mitochondria following glabridin treatment occurred substantially earlier than the swelling of the endoplasmic reticulum. Based on the above results, we hypothesize that the reduction in cell activity caused by glabridin is directly connected to mitochondrial dysfunction.

Mitochondrial membrane potential (MMP) is essential for the respiratory chain generating ATP, and its absence causes mitochondrial dysfunction and the release of reactive oxygen species (ROS) [33]. A previous report indicated that glabridin promotes oxidative stress by producing reactive oxygen species (ROS) and induces MMP depolarization [34]. Similarly, our findings suggest that glabridin treatment promotes MMP loss as well as enhanced ROS formation. However, the involvement of ROS generation and MMP loss in glabridin-induced paraptosis-like cell death has to be investigated further.

In conclusion, our findings reveal that glabridin enhances cytoplasmic vacuolation as a result of increased ER stress and proteasome suppression, leading to paraptosis-like cell death. Glabridin also caused mitochondrial dilatation, MMP loss, and ROS production, all of which contributed to this process. The molecular processes underlying glabridin-induced paraptosis-like cell death are being investigated further.

## Declarations

## Author contribution statement

X.C. and M.C. designed and performed the experiments, analyzed and interpreted the data, and wrote the manuscript. M.C. is responsible for the execution of the project, data analysis, and manuscript preparation.

## Data availability statement

Data will be made available on request.

