## supplemental information for "Glabridin induces paraptosis-like cell death via ER stress in breast cancer cells"

### Supplementary Information

### Figure S1.

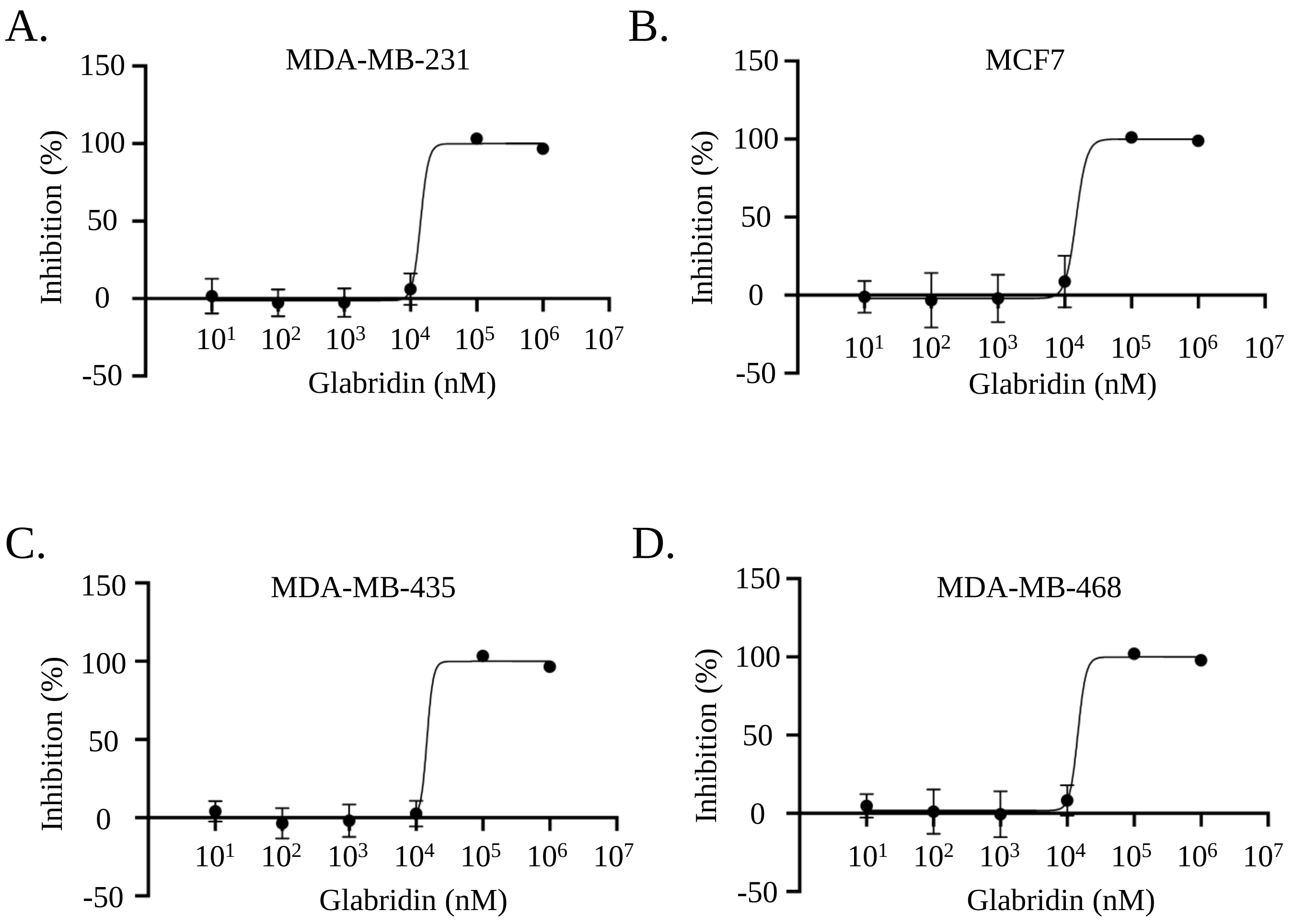


**Figure S1.** Dose response inhibition curves of glabridin on human breast cancer cell lines. (A) MDA-MB-231 cells, (B) MCF7 cells, (C) MDA-MB-435 cells, (D) MDA-MB-468 cells. Cells were treated with glabridin at the indicated concentrations for 24 h, and cell proliferation inhibition rate was determined by CCK-8 assay. All error bars were represented in mean ± SE.

### Figure S2.

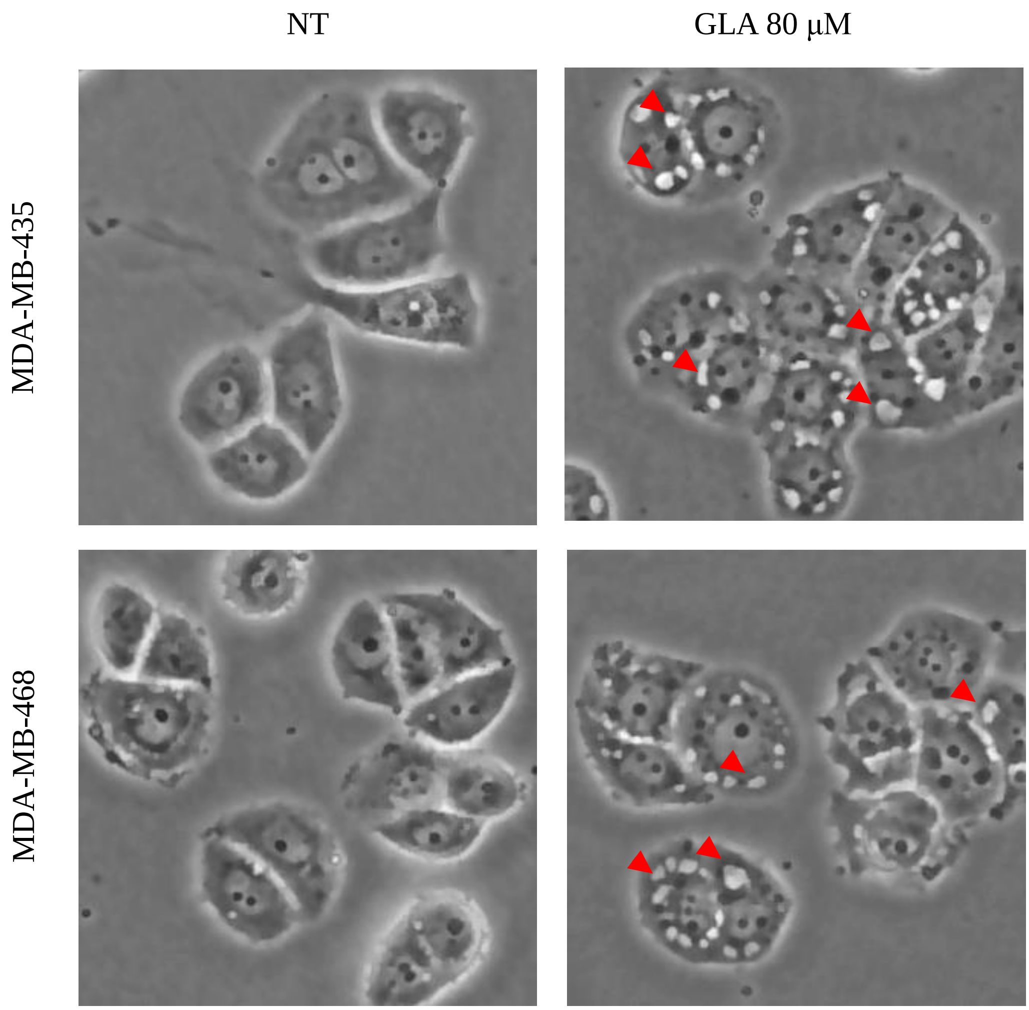


**Figure S2.** Morphological changes of MDA-MB-435 and MDA-MB-468 cells after 6 h treatment with 80μM glabridin observed under an inverted light microscope. Cytoplasmic vacuoles were indicated by the red arrows.

### Figure S3.

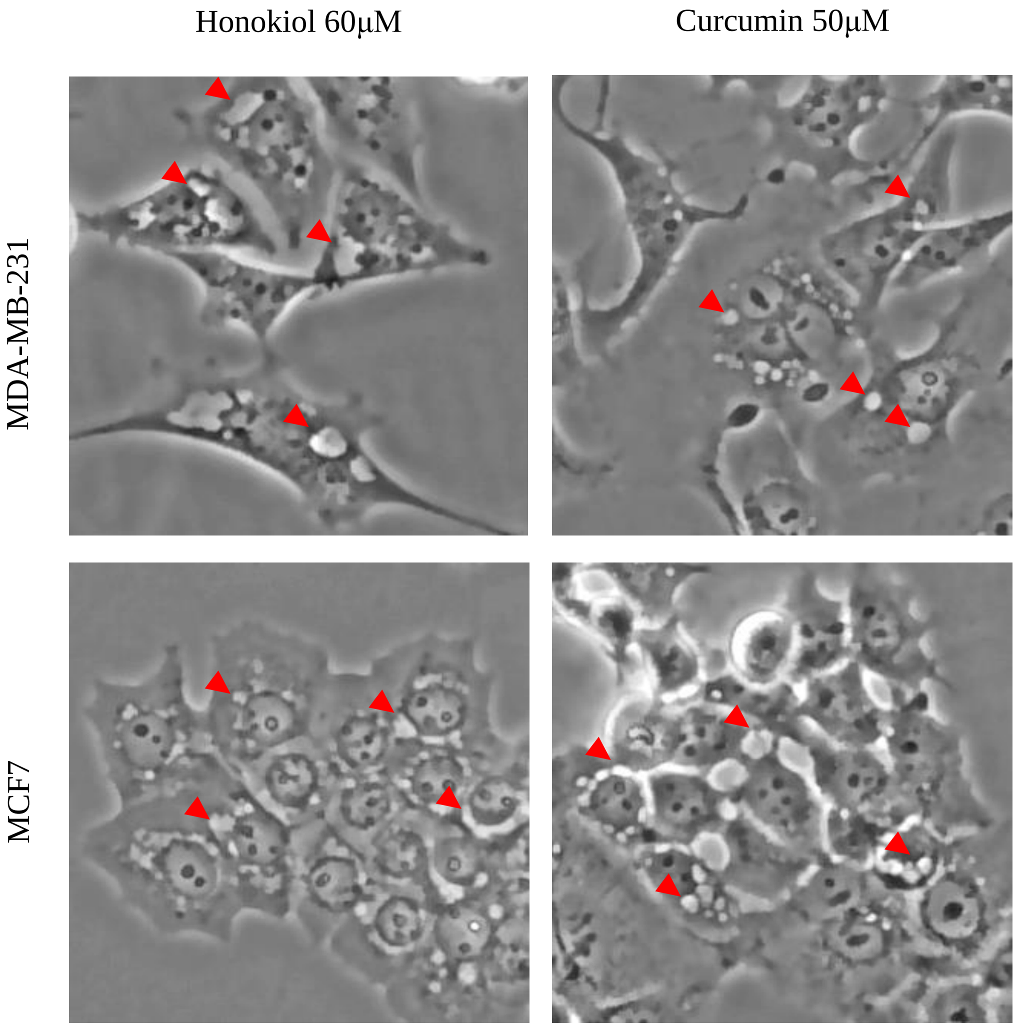


**Figure S3.** Inverted light microscopy images of MDA-MB-231 and MCF7 cells after 6 h of treatment with 60μM honokiol or 50μM curcumin. The red arrows pointed to cytoplasmic vacuoles.

### Figure S4.

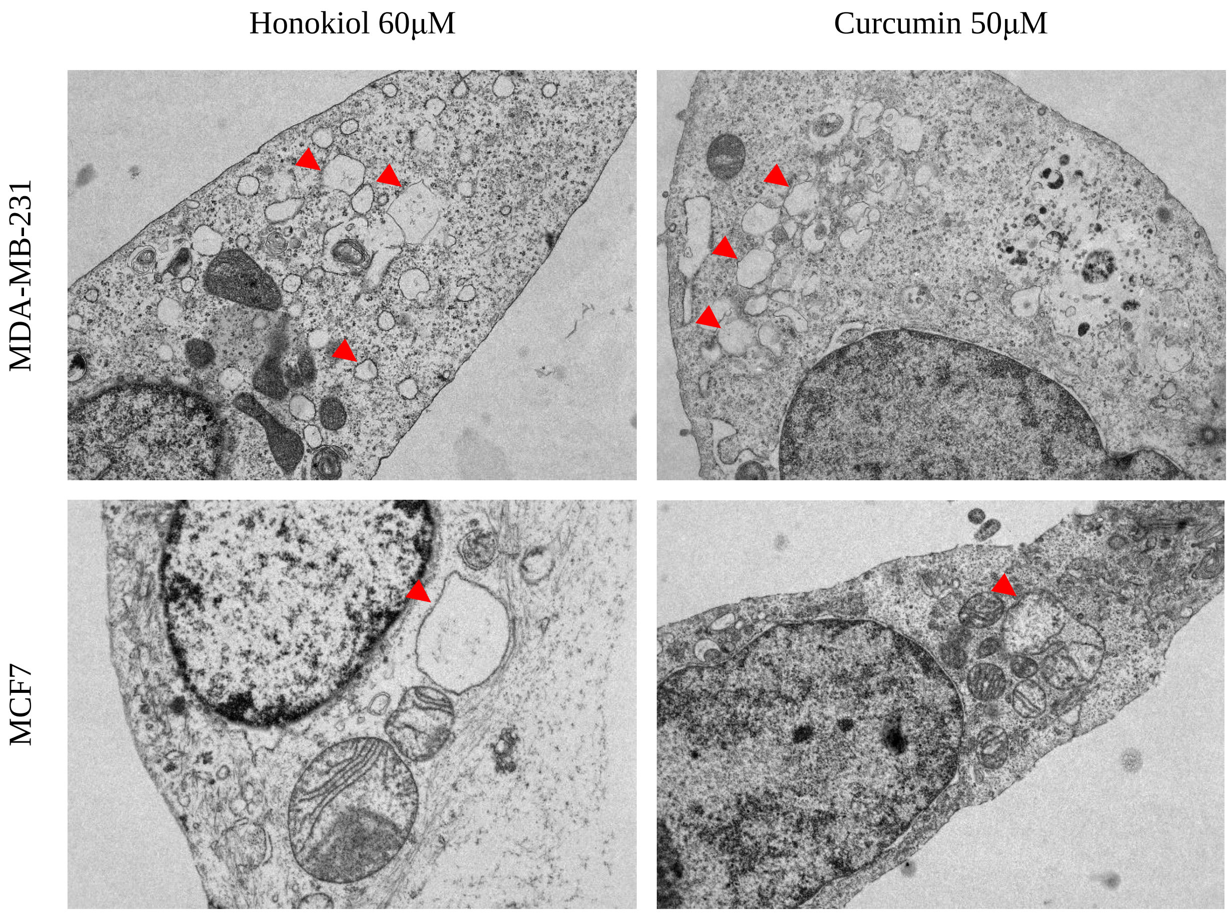


**Figure S4.** Transmission electron micrograph images of cells treated with honokiol (60 μM) or curcumin (50μM) for 6 h. Cytoplasmic vacuoles were indicated by the red arrows.

### Figure S5.

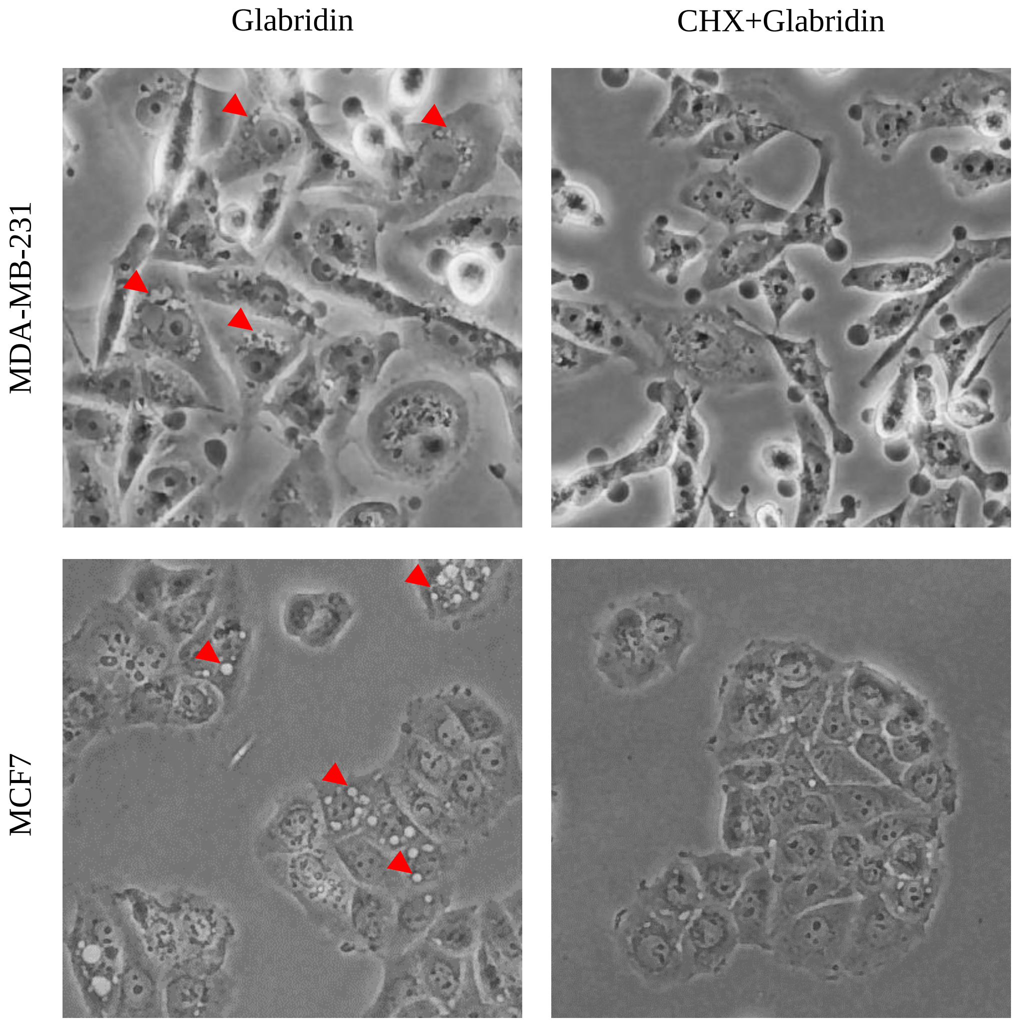


**Figure S5.** Representative images of MDA-MB-231 and MCF7 cells co-treated with glabridin and cycloheximide. Cells were treated with 80µM glabridin for 3.5h in the absence and presence of 500nM CHX, and fixed with 4% paraformaldehyde for 30 min at room temperature. Cytoplasmic vacuoles were indicated by the red arrows.

### Figure S6.

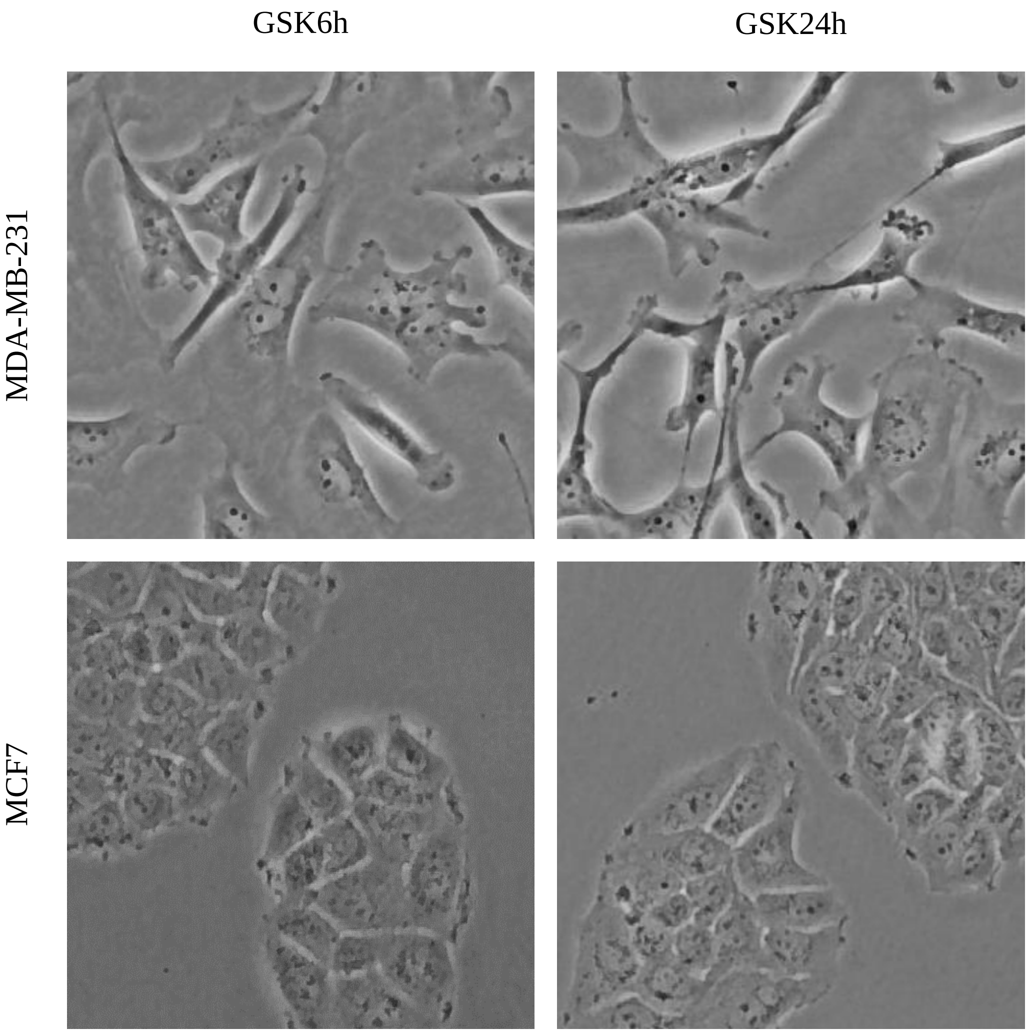


**Figure S6.** Inverted light microscopy images of MDA-MB-231 and MCF7 cells treated with 1μM GSK2656157 for indicated times.

Table S1.

| Gene | Forward primer | Reverse primer |
|---|---|---|
| BiP | 5’-GTTTGCTGAGGAAGACAAAAAGCTC-3’ | 5’-CACTTCCATAGAGTTTGCTGATAAT-3’ |
| CHOP | 5’-GGAAACAGAGTGGTCATTCCC-3’ | 5’-CTGCTTGAGCCGTTCATTCTC-3’ |
| XBP-1 | 5’-AGCCAAGGGGAATGAAGTG-3’ | 5’-CTGAAGAGTCAATACCGCC-3’ |
| β-actin | 5’-TCCTCCCTGGAGAAGAGCTAC-3’ | 5’-TCCTGCTTGCTGATCCACAT-3’ |
